# Investigation of *Aspergillus fumigatus* small RNA biogenesis uncovers evidence of double-stranded RNA-dependent growth arrest

**DOI:** 10.1101/2024.05.24.595671

**Authors:** Abdulrahman A. Kelani, Xiaoqing Pan, Swatika Prabakar, Lukas Schrettenbrunner, Matthew G. Blango

## Abstract

*Aspergillus fumigatus* is a ubiquitous filamentous fungus and dangerous human pathogen that produces a limited pool of small RNAs, consisting in large part of tRNA-derived RNAs (tDRs). Here, we improve our understanding of the small RNAs produced in conidia and mycelium of *A. fumigatus* using small RNA-sequencing of strains lacking RNA interference (RNAi) machinery and a cutting-edge tDR-sequencing approach. We find little evidence of small RNAs dependent on the canonical RNAi machinery under laboratory growth conditions, but reveal tDRs to be differentially abundant across fungal morphotypes, with specific fragments proving dominant in each assessed condition (e.g., Asp(GTC)-5’tRH in conidia; His(GTG)-5’tRH in mycelium). Consistent with the literature, we observed distinct patterns of tDRs from nuclear- and mitochondria-derived tDRs, which was confirmed for wild-type fungus with tDR-seq. By inducing canonical RNAi with overexpression of an inverted-repeat transgene, we determined *A. fumigatus* to produce predominantly 20-nt, 5’ uridine-containing small RNAs from the transgene, a population reliant on Argonaute and Dicer-like proteins. Surprisingly, we found that overexpression of this double-stranded RNA (dsRNA) limited growth of both wild-type and RNAi-deficient strains, with a strain lacking the two *A. fumigatus* RNA-dependent RNA polymerase orthologs particularly vulnerable. Dicer-like- and Argonaute-double knockout strains with few detectable transgene-derived small RNAs were also susceptible to growth inhibition, suggesting excessive dsRNA limits growth in *A. fumigatus*. Ultimately, we have provided an improved description of small RNA biogenesis in *A. fumigatus* and uncovered an intriguing link between dsRNA metabolism and filamentous growth.

**IMPORTANCE:** The mechanisms of small RNA biogenesis in fungi are diverse and multifactorial. Here, using a panel of RNA interference gene deletion strains and multiple small RNA sequencing approaches, we reveal limited input of the canonical RNAi machinery on the small RNA landscape of the human fungal pathogen, *Aspergillus fumigatus* under normal growth conditions. We identify specific patterns of tRNA-derived RNAs unique to fungal morphotypes, which may hold promise for future diagnostic efforts. We then probed the mechanism of double-stranded RNA processing by the organism after artificial induction, revealing a potential link between proper double-stranded RNA processing and fungal growth arrest, improving our understanding of the evolution of double-stranded RNA regulation in a ubiquitous mold.

## INTRODUCTION

The canonical RNA interference (RNAi) pathway processes double-stranded RNA (dsRNA) through cleavage to small RNAs (sRNA) that are then used as guide molecules to facilitate silencing of target transcripts (1). The dsRNA precursor molecule can be produced endogenously via transcription or exogenously, e.g., from a virus or repetitive transgene. The dsRNA A-form helix is characterized by a small major groove, unlike that of B-form DNA, meaning that the molecule itself can be a substrate for multiple dsRNA binding proteins (dsRBPs) independent of the nucleotide sequence (2). The result is competition for dsRNA substrates by dsRBPs that bind, cleave, or react to the dsRNA molecule through interactions with the sugar-phosphate backbone (2). In higher eukaryotes, this manifests as a balance between processing (RNA cleavage or modification) and activation of inflammatory cascades by dsRBP sensors like MDA5 (3), whereas in microbes like the bacteria *Escherichia coli*, dsRNA is thought to be rapidly degraded or processed to smaller RNAs (4).

The sRNA landscape of fungi (like many eukaryotes) consists of a heterogenous group of RNA molecules, ranging from fragments of housekeeping RNA like tRNA and rRNA to small interfering RNAs (siRNA) and microRNA-like RNAs (milRNA) produced from both perfectly duplexed dsRNA and dsRNA structures with bulges and/or mismatches (5). Fungal sRNAs are produced from these precursors via diverse biogenesis pathways, often including members of the canonical RNAi machinery such as Dicer (or Dicer-like), Argonaute, and/or RNA-dependent RNA polymerases (RdRPs), together with peripheral factors (6). Some fungi have lost RNAi to gain an advantage in a particular niche (7), but many, including the ubiquitous human pathogen Aspergillus fumigatus, harbor an active RNAi pathway (8). Interestingly, *A. fumigatus* appears to produce a limited pool of endogenous sRNAs under standard growth conditions, consisting in large part of tRNA-derived RNAs (tDRs) despite encoding two Dicer-like proteins, two Argonautes, and two RdRPs (9-11). The *A. fumigatus* RNAi system is inducible in fungal mycelium and capable of processing dsRNA from inverted-repeat transgenes (IRT) and mycoviruses (8, 9), but the alternative functions of RNAi like tDR biogenesis remains unclear.

The tDRs are a heterogeneous group of 10-to 50-nucleotide (nt) sRNAs that include the 5’ and 3’ tRNA halves (tRHs), tRNA fragments (tRF) processed from the 5’ (tRF-5) and 3’ ends (tRF-3, tRF-3 CCA), and internal tRFs (i-tRF) (12) (**Fig. S1**). The generation of tRHs (∼31 to 40 nt) is multifactorial, but cellular stress is a well-described trigger of tRNA cleavage particularly in the anticodon loop, resulting in tRNA-derived, stress-induced fragments (tiRNAs) (11, 13-15). Enzymes like the RNase A ribonuclease angiogenin and other endonucleases like RNase T2 cleave tRNAs in the anticodon loop (16, 17). tRF-5 and tRF-3 are 17 to 25 nt long products of mature tRNA cleavage at the 5⍰ end D loop and 3’ end T loop, and are produced by angiogenin or the RNase III enzyme Dicer in some species (18-21). Interestingly, the biogenesis of tRFs and tRHs is dynamic across development (11, 22). In *A. fumigatus*, tRHs accumulate after germination from conidia to hyphae independently of mature tRNA levels; however, during the early conidiogenesis there was a rapid increase in tRHs and a decrease in mature tRNA levels (11). These results suggested that the decrease in the mature tRNA pool in conidia may contribute to the metabolic dormancy of spores as active translation is shut down.

tRNA cleavage is a conserved eukaryotic response to not just starvation but also other stress conditions, e.g., oxidative stress induces tiRNA production in both mammalian cells and Saccharomyces cerevisiae (15, 25). In yeast, endonucleolytic cleavage of tRNA is performed by Rny1p, an ortholog of RNase T2 (28). Several other stress conditions such as heat stress, UV stress, glucose starvation, nitrogen starvation, and amino acid starvation all resulted in differing patterns of tRNA cleavage in S. cerevisiae (15). In *Tetrahymena thermophila*, it was shown that tRF-3 RNAs associate with the Argonaute/Piwi protein, Twi12, and promote its translocation into the nucleus to facilitate ribosomal RNA processing (29). tRF-3 fragments that interact with the Argonaute-containing RNA-induced silencing complex (RISC) appear able to guide silencing of target mRNA based on 3’UTR seed sequence complementarity (28, 30). The RNAi pathway is known to use sRNAs or even tRFs as guides to induce both transcriptional gene silencing of target gene loci and post-transcriptional gene silencing of target RNA transcripts, suggesting complex interplay between the machinery and substrates.

We previously showed that overexpression of a long dsRNA in the form of an inverted-repeat transgene resulted both in silencing of a nonessential target gene with sequence complementarity and in decreased fungal biomass (31). These results are interesting in the context of the varied growth effects reported after infection of *Aspergillus species* with mycoviruses (32-35), and in light of our observed contribution of the RNAi system to regulation of expression of ribosome biogenesis genes in *A. fumigatus* conidia (9). These findings led us to hypothesize that overproduction of long dsRNA substrates in *A. fumigatus* sequesters dsRBPs away from housekeeping tasks, potentially limiting growth. In fact, Dicer proteins have many functions in addition to their canonical role in creating sRNAs (36). In humans, a lone Dicer, DICER1, associates with RNA polymerase II in the nucleus to prevent accumulation of dsRNA during active transcription and limit unwanted activation of interferon pathways by unintentional production of dsRNA (37, 38). These interferon pathways are well-integrated with Dicer biology in higher eukaryotes, whereas more evolutionarily ancient Dicers have a more direct role in the antiviral response reliant on the ATPase activity of their helicase domains, an activity lost in humans (39). Although mycoviruses cause varied responses, our understanding remains rudimentary as to how lower eukaryotes like fungi that pre-date the interferon response control and respond to excess dsRNA.

Here, we provide an improved global characterization of the sRNA landscape of the World Health Organization critical priority pathogen *A. fumigatus* (40) using sRNA-seq and tDR-seq. We reveal differential abundance of tDRs from *A. fumigatus* conidia (asexual spores) and mycelium (hyphal network). We go on to show that heterologous overexpression of an inverted-repeat transgene results in production of small RNAs and limits fungal growth even in the absence of a functional RNAi system, suggesting that excess dsRNA may itself serve as a check on growth in *A. fumigatus*.

## RESULTS

### A. fumigatus encodes five annotated proteins with putative RNase III activity

Many sRNAs are generated from structured precursors by cellular dsRBPs with RNase III domains capable of cleaving dsRNA. *A. fumigatus* has 10 annotated dsRBPs (FungiDB.org), with 5 being putative RNase III enzymes (InterPro). These contain two Dicer-like proteins (DclA, Afu5g11790; DclB, Afu4g02930), a mitochondrial RNase III enzyme that is an ortholog of MRPL3 from N. crassa (Afu1g12730; (6)), the nucleolar RNase III enzyme Rnt1 (Afu5g04440), and a small RNase III domain-containing protein of unknown function found only in *A. fumigatus* and a few close relatives (Afu5g06830) (**Fig. S2A**). The five other dsRBPs are less well-studied, but in several cases have roles in rRNA assembly or function (Ribosomal Protein S5, Afu7g01460; 37S Ribosomal Protein S5, Afu5g11540; RAD52 ortholog, Afu4g06970; Afu2g16140; and Afu4g10320). Two of these dsRBPs are predicted to reside in the mitochondria (MRPL3; 37S Ribosomal Protein S5), with the others likely found in the cytoplasm or nucleus. Expression profiling from previously published mRNA-seq data on conidia, 24-h mycelium, and 48-h mycelium revealed stage-specific expression of many of these genes often influenced by expression of the core RNAi components ((9) and Fig. S2B), suggesting that specific subsets of proteins might compete for dsRNA binding at different growth stages. As deletion of the Dicer-like proteins resulted in dysregulation of the conidial transcriptome in our previous work (9), we were interested to define the Dcl-dependent sRNAs of *A. fumigatus*.

### The composition of A. fumigatus endogenous sRNAs is morphotype specific

Using previously described knockout strains in the canonical RNA interference machinery (*ΔdclA/B, ΔppdA/B*, and *ΔrrpA/B*) created in the wild-type CEA17Δ*akuB*^KU80^ background, we simultaneously profiled expression of small non-coding RNA (ncRNA; this study) and mRNA (9) from the same biological samples across three conditions (conidia, 24-h mycelium, and 48-h mycelium; (**Fig. 1A; Fig. S3A; Dataset S1**)). We mapped the expression of distinct small ncRNA classes, including snRNA, snoRNA, and tDRs (**Fig. 1B**). Across all the sequencing samples, rRNA made up the majority of the mapped reads, followed by short reads mapping to tRNA genes. In conidia, many reads mapped to rRNA, whereas the proportion of tDRs and other small ncRNAs increased in the mycelial samples, consistent with an expected, more diverse transcriptome in metabolically active mycelium. The regulatory potential of these fragments remains poorly understood, particularly in filamentous fungi.

**FIG 1.**
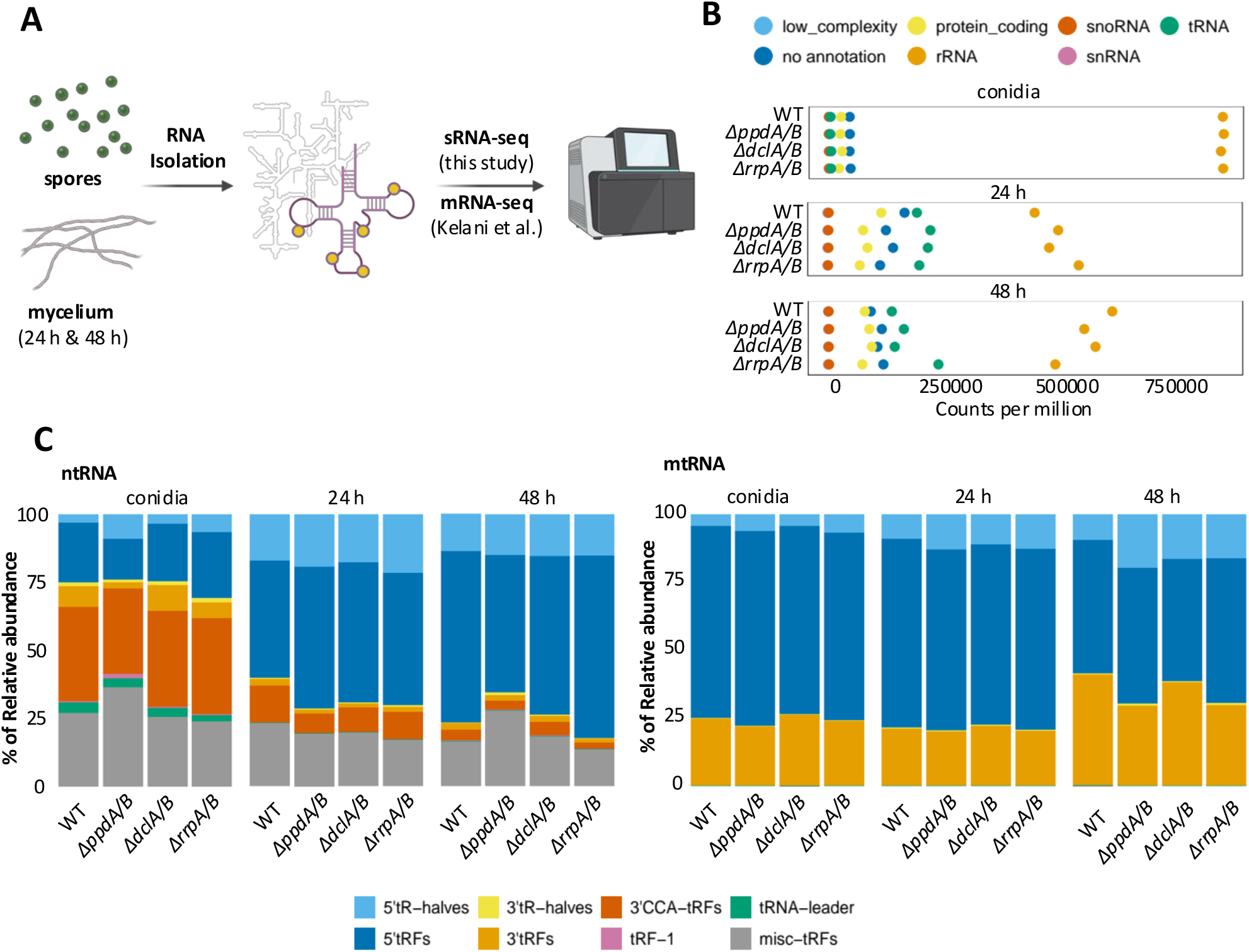
Small RNA abundance in A. fumigatus conidia and mycelium. (**A**) Strains and conditions profiled in this study. The wild-type (CEA17 ΔakuB^ku80^) and three knockout strains (Δ*ppdA*/B, Δ*dclA*/B, Δ*rrpA*/B) were collected (n = 3 biological replicates per condition). RNA samples were collected from fungal conidia and hyphae (24 hours and 48 hours) and Illumina sequenced. Created in BioRender. Blango, M. (2025) https://BioRender.com/r33p545. (**B**) Coverage of small ncRNA types within the profiled samples. Small ncRNAs were annotated with low complexity, no annotation, protein coding, rRNA, snoRNA, snRNA, and tRNA (including nuclear tRNA and mitochondrial tRNA). (**C**) Comparison of tDR relative abundance across samples, and between wild-type and RNAi double knockouts for both nuclear-(ntRNA) and mitochondria-encoded tRNA (mtRNA). n = 3 biological replicates per condition.

We next aimed to identify sRNAs that might be masked by the high abundance of rRNA reads. Removal of reads matching rRNA and restriction of read length to 18 to 30 nt led to identification of similar numbers of sRNAs from each of the double knockout strains compared to wild type (**Fig. S3B-C**). In conidia, a clear population of 18 to 21 nucleotides was identified, whereas in 24-h and 48-h mycelium, we generally observed two or more peaks centered at ∼19 nt and/or ∼28 nt, often indicative of tRNA-derived RNAs. We then used Manatee (41) to identify and predict sRNAs, including microRNA-like RNAs (milRNA: (**Dataset S2**)). We observed few low abundance candidate sRNAs in conidia, and a larger number of slightly higher abundance sRNAs from 24-h and 48-h mycelium. Interestingly, deletion of *dclA*/B did not change the overall number of candidate sRNAs identified (1,524 compared to 1,615 in wild type), whereas loss of *ppdA*/B and *rrpA*/B decreased the number of sRNAs identified by Manatee (873 and 1,016 respectively), although most of the candidates identified were not of particularly high abundance (**Dataset S2**). These results are consistent with studies showing a limited pool of sRNAs meeting the criteria of milRNA in *A. fumigatus* (9-11), and instead more RNAs likely derived from mRNA turnover or structural RNAs (e.g., tRNAs and rRNA).

### tDR biogenesis is primarily independent of the RNAi machinery in A. fumigatus

*A. fumigatus* produces an abundance of tRHs during starvation (11). In agreement we observed an increase in the relative abundance of nuclear-encoded 5’-tRHs from conidia to older mycelium; however, we also detected a large abundance of tRFs, likely generated by cleavage of either mature or precursor tRNA (e.g., by RNase T2). For nuclear-encoded tRFs, we noticed that the relative abundance of tRF-5 increases from conidia to 24-h-old and 48-h-old mycelium where nutrients are more limited, while the tRF-3 abundance decreased across those samples (**Fig. 1C**). However, within mitochondrial tDRs, reads mapped mostly to tRF-3, 5’-tRHs, and tRF-5 (**Fig. 1C**).

Next, we created normalized heat maps of abundance for the tDRs of each tRNA (**Fig. 2; Fig. S4**) using the unitas software package as previously described (42). We observed clear differences in the relative abundance of tDRs from each morphotype, suggesting differential stability or potentially poor ligation following cleavage. We saw slight differences in tDR biogenesis for some of the RNAi knockout strains, especially Δ*ppdA*/B, as compared to the wild type, but according to Benjamini-Hochberg-corrected P-values determined by t test, these were not significantly different. Nonetheless, in conidia, Δ*ppdA*/B appeared to produce fewer tRF-3, tRF-3 CCA, and tRF-5 derivatives from many tRNAs. Surprisingly, we saw minimal differences in the production of tDRs in Δ*dclA*/B, with most of the differences occurring for tRF-5 at 48 h. These results suggest that the RNAi machinery, particularly the Dicer-like proteins, play only a minor role in tRNA fragment production in *A. fumigatus* under standard growth conditions.

**FIG 2.**
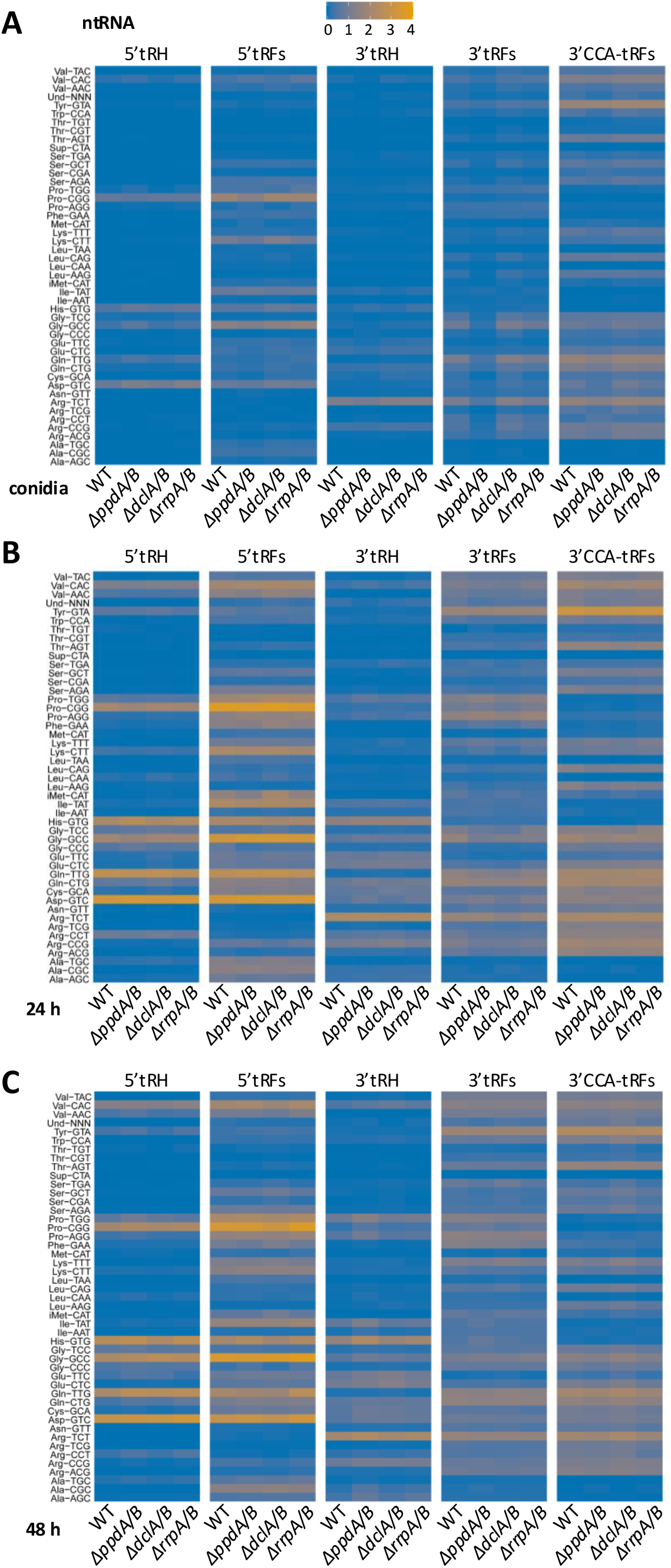
Expression of tDRs in A. fumigatus conidia and mycelium. Heatmap showing abundance of tDR species derived from nuclear-encoded tRNA (ntRNA; tRNA Transcript ID =1, Gene locus ID =1) across sampled morphotypes: (**A**) Conidia, (**B**) 24-h-, and (**C**) 48-hold-mycelium of wild-type and RNAi double knockouts as determined by sRNA-seq. Plots show counts per million mapped reads based on fractionated counts produced with unitas v. 1.7.0 (55). N = 3 biological replicates per condition.

### tDR-seq reveals altered patterns across fungal morphotypes

The abundance of tDRs in our sRNA-seq together with the known difficulties of sequencing tRNAs using standard methodologies (e.g., poor adapter ligation and decreased reverse transcription efficiency) led us to perform a tRF & tiRNA-seq (here denoted tDR-seq) approach, where RNA ends are repaired and common methylation modifications are removed to facilitate ligation and readthrough, respectively (**Fig. 3A; Fig. S5; Dataset S3**). We focused on the wild-type fungal morphotypes for simplicity as no major differences were observed between the RNAi knockouts and wild type in the previous analysis. In tDR-seq, more reads mapped to rRNA fragments than in the sRNA-seq; however, a closer look at the distribution of tDRs indicated a higher proportion of tRHs compared to the sRNA-seq analysis for both the nuclear-encoded and mitochondria-encoded tRNAs (**Fig. 3B**). As compared to the tDR distribution from sRNA-seq (**Fig. 1C**), tDR-seq revealed more 3’tRHs relative to other tDRs, consistent with an approach that fixes RNA ends prior to library preparation (**Fig. 3B**). Interestingly, the relative abundance of the 5’tRHs with nuclear origin were much higher in the conidia (∼75%) after the tDR-seq approach (**Fig. 3B**). Further analysis revealed that a majority (∼95%) of tDRs identified in the conidia came from a particular Asp(GTC) nuclear-encoded tRNA, with the 5’tRH being the most abundant fragment (**Dataset S3**). The length of the tRNA fragments was more consistent with tRHs for the nuclear-encoded tRNAs, but the mitochondria-encoded tDRs had a wider range, with a particular relative enrichment of smaller fragments in hyphae (**Fig. 3C**), consistent with the literature (42).

**FIG 3.**
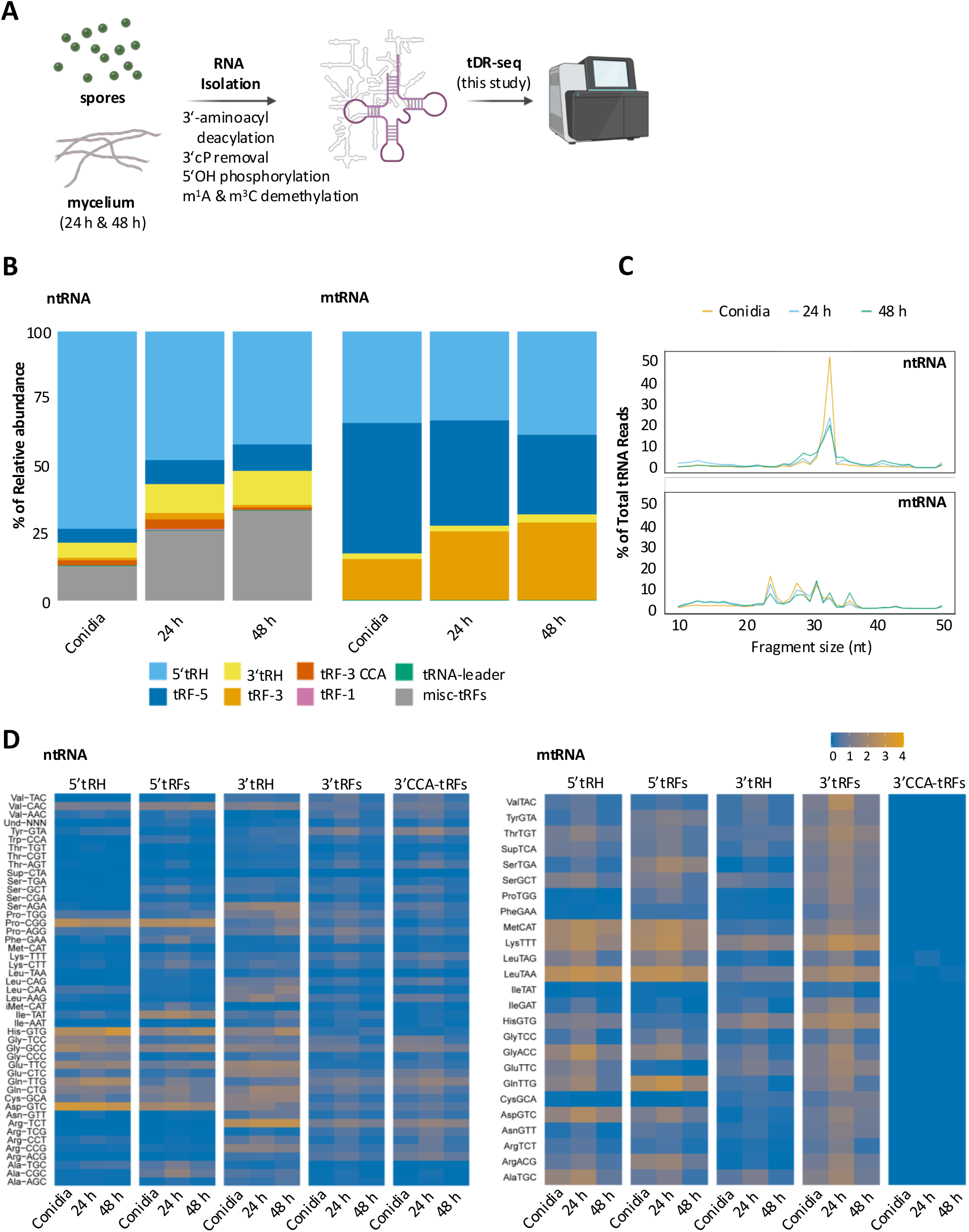
Dynamics of tDR expression in A. fumigatus conidia and mycelium revealed by tDR-seq. (**A**) RNA samples (n = 3) were collected from wild-type *A. fumigatus* conidia and hyphae. Samples were pre-treated to remove specific modifications and resolve tDR ends prior to library preparation. RNA molecules of up to 50 nt were selected for tDR-seq using Illumina sequencing. Created in BioRender. Blango, M. (2025) https://BioRender.com/e54f907. (**B**) Relative abundance of different tDRs across samples for tDRs derived from nuclear- (ntRNA) and mitochondria-encoded (mtRNA) tRNAs. (**C**) Average tDR length identified across samples for tDRs derived from either ntRNA (**top**) or mtRNA (**bottom**). (**D**) Heatmap showing abundance of nuclear- (ntRNA) and mitochondria-encoded (mtRNA) tDR species and their tRNA of origin (tRNA Transcript ID =1, Gene locus ID =1) across sampled conidia and mycelium timepoints from tDR-seq dataset. Plots show counts per million mapped reads based on fractionated counts produced with unitas v. 1.7.0 (55). n = 3 biological replicates per condition.

A heat map presenting the tDRs for each tRNA again revealed substantial differences in abundance and fragment distribution between tRNAs with different or even the same isoacceptor (**Fig. 3D**). Even though many of the detected tRHs derive from Asp(GTC), other tRNAs like His(GTG) and Pro(CGG) also produce a relatively high abundance of tRHs. Many of the nuclear-encoded tRNAs appeared to have stable tRF-3 CCA fragments, with the previously identified Tyr(GTA)-tRF-3 CCA being among the most abundant fragments (**Fig. 3D**; (9)). Curiously, tRF-3 CCA were almost completely lacking from mitochondria-encoded tRNAs (**Fig. 3D; Fig. S4A-C**), with a closer investigation only revealing a few poorly mapped reads. Further work will be required to experimentally validate this result. Collectively, these results corroborate that the *A. fumigatus* small transcriptome consists mostly of sRNAs derived from structural RNAs (e.g., rRNAs and tRNAs) under standard growth conditions.

### The A. fumigatus RNAi machinery processes inverted-repeat transgene dsRNA to 20-nt, 5’U-containing sRNAs

*A. fumigatus* is known to produce sRNAs during mycovirus infection, but in this case, the identified RNAs lacked canonical fungal sRNA properties (e.g., a 5’U) (10). To assess the sRNA environment in a more defined system, we turned to our previously used *pksP* IRT system (9), which in liquid culture led to a surprising growth inhibition of *A. fumigatus* (43). We initially postulated that the inhibition phenotype was reliant on off-target effects introduced by production of sRNAs from our 500-bp *pksP* IRT with unanticipated complementarity to essential genes. To further test this hypothesis, we produced *pksP* IRT-expressing RNAi knockout strains (Δ*dclA*/B-IRT, Δ*ppdA*/B-IRT, and Δ*rrpA*/B-IRT) and induced expression of the transgene in liquid culture. We grew the *pksP* IRT strains for 16 h plus 8 h of induction to ensure sufficient biomass prior to collection of RNA and sRNA-seq analysis (**Fig. S6A, Dataset S4**). This allowed us to assess the ability of *A. fumigatus* to produce sRNAs from the transgene transcript. Untreated wild-type fungus was included as a control for standardization with previous experiments. sRNAs were mapped to the *pksP* IRT sequence, and we observed an accumulation of 20-nt RNAs with a 5’uridine produced from the transgene in the wild-type and Δ*rrpA*/B-IRT strains, revealing that the *A. fumigatus* RNAi machinery can function in a manner consistent with the fungal literature (**Fig. 4A-C**). These peaks of sRNA were absent in the wild-type, Δ*dclA*/B-IRT, and Δ*ppdA*/B-IRT strains. Closer inspection of the mapped reads revealed the largest accumulation of mapped sRNA reads originated from the 3’half of the transcript (antisense) primarily as approximately two 20-nt RNAs (**Fig. 4C**). Prediction of target binding sites using these two relatively abundant sRNA sequences revealed no additional high-quality targets beyond the complementarity to the *pksP* gene, potentially suggesting that no or other less abundant sRNAs facilitate the toxicity observed by the expression of the inverted-repeat transgene in our previous study (9).

**FIG 4.**
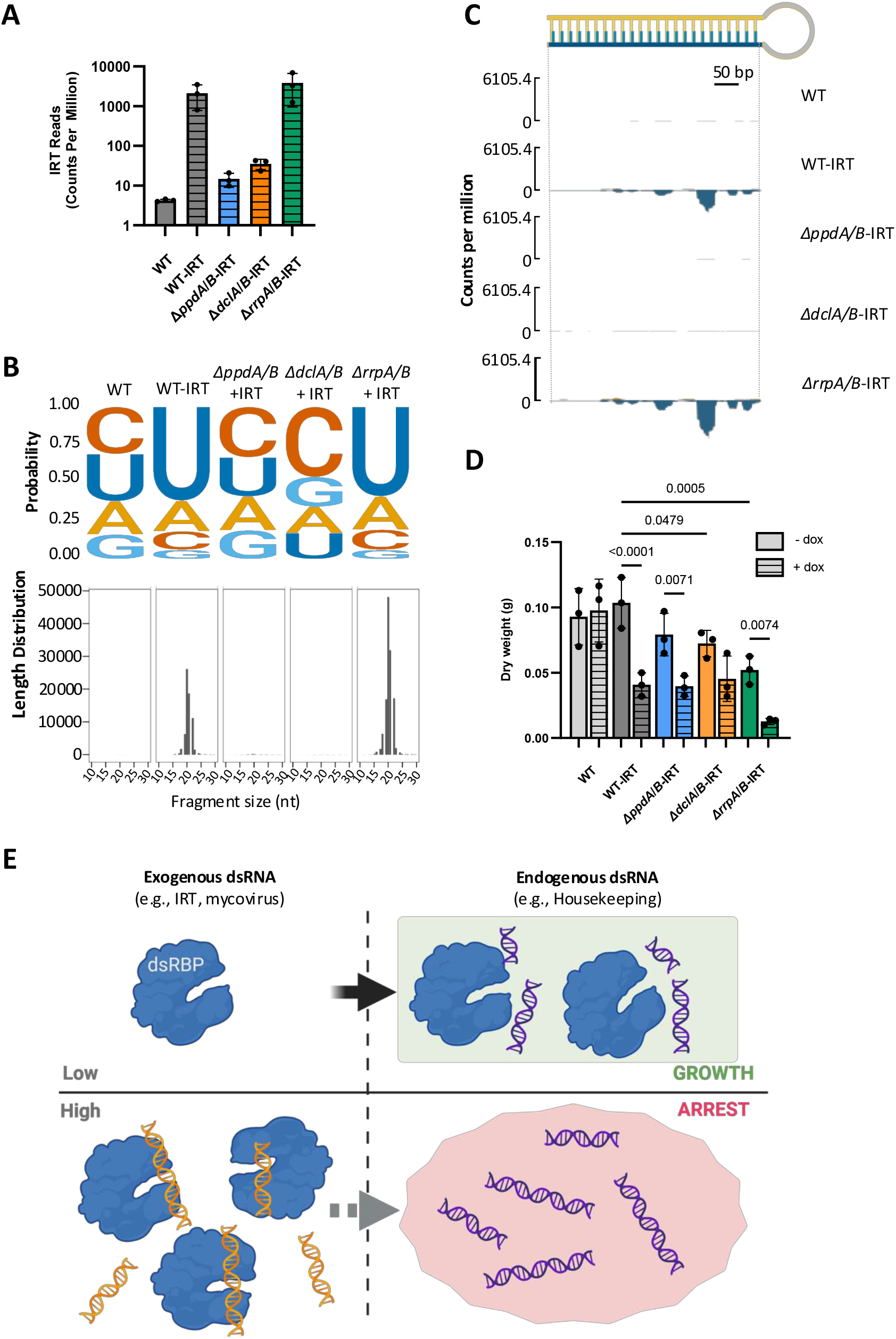
Inverted-repeat transgene expression induces 20-nt, 5’U-containing sRNA production. (**A**) sRNA reads mapped to *pksP* inverted-repeat transgene sequence per million mapped reads for each of the denoted strains. Data is accumulated from three biological replicate sRNA-seq experiments. (**B**) The distribution of the 5’ nucleotide for the respective strains (**top**) and histogram showing the average sRNA read length for the sRNAs that mapped to the *pksP* inverted-repeat transgene (**bottom**). (**C**) Graph showing read coverage of *pksP* inverted-repeat transgene region created with SparK package (59) with reads from the 5’ end of the inverted repeat shown as positive strand and from 3’ end shown as negative strand. (**D**) Measurement of fungal dry weight (biomass) after 24-h growth in liquid culture in the presence or absence of doxycycline as an inducer of *pksP* IRT in the RNAi knockout strains or compared to a wild-type strain. Relevant comparisons were performed with two-way ANOVA and Šídák’s multiple comparisons posttest from three biological replicates. (**E**) Working model of dsRNA-induced growth arrest in *A. fumigatus*. High levels of exogenous dsRNA sequester dsRBPs from normal function and limit growth rate. Created in BioRender. Blango, M. (2025) https://BioRender.com/e82n319.

### dsRNA overexpression limits A. fumigatus growth

We next tested the contribution of sRNA production to the observed growth inhibitory effect of the *pksP* IRT in liquid culture. As expected, induction of the inverted-repeat transgene in the wild-type strain limited fungal biomass as before (9), and this phenotype after induction was largely diminished in the Δ*dclA*/B strain (**Fig. 4D**). Low expression of dsRNA via leaky TetON induction in each of the knockout strains resulted in decreased growth even in the uninduced RNAi double knockout strains, suggesting that dsRNA may itself be growth inhibitory. Interestingly, the Δ*ppdA*/B and Δ*rrpA*/B strains continued to show significant growth decreases in biomass after induction. The Δ*rrpA*/B double knockout was particularly susceptible to overexpression of dsRNA stress in this experiment. As a control, we grew each of these strains in the absence of the inverted-repeat transgene, with no major differences in growth observed for the RNAi double knockout strains (Fig. S5B). Finally, we performed a similar unitas analysis on our IRT-sRNA-seq experiment to assess the influence of dsRNA overexpression on tDR biogenesis. We found that the wild-type and Δ*rrpA*/B strains harboring the *pksP* IRT exhibited slight increases in tRH abundance and decreases in tRFs compared to the Δ*dclA*/B and Δ*ppdA*/B strains (**Fig. S6C**). This trend was noticeable for tDRs from both the nuclear and mitochondria-encoded tRNAs and implies a potential link between dsRNA metabolism and tRNA processing in *A. fumigatus*.

## DISCUSSION

Small ncRNAs are key regulators of gene expression and stress response in many organisms, including fungi. Here, we provide an improved description of the sRNA landscape of *A. fumigatus* by leveraging sRNA-seq and tDR-seq. The pool of *A. fumigatus* tDRs appears to be heterogeneous, dynamic, and only partially dependent on the canonical RNAi machinery under standard growth conditions. Canonical 20-nt 5’-uridine-containing sRNAs can be produced from *A. fumigatus* mycelium upon induction of an inverted-repeat transgene. Expression of dsRNA had the simultaneous effect of limiting fungal growth in an RNAi-independent manner.

We initially performed sRNA-seq to evaluate the dynamics of small ncRNAs during two distinct morphological phases of *A. fumigatus* growth and identified expected classes of small ncRNAs, with rRNA and tDRs exhibiting the highest abundance in conidia and mycelium. Although often a sign of RNA degradation, our experience with *A. fumigatus*, in combination with our quality control steps and the literature (11), suggests that rRNA and tRNA-derived RNAs do accumulate over the lifecycle, particularly at later growth timepoints (e.g., 48-h mycelium). Interestingly, the abundance of tDRs observed in our sRNA-seq and tDR-seq did not appear to correlate with our previous measurement of the overall tRNA abundances from mycelium under the same growth conditions using Nano-tRNAseq, where for example, Ser(GCT) was found to be the most abundant tRNA isoacceptor (44). This supports the hypothesis that tDR biogenesis is a regulated process in *A. fumigatus*, and not solely a product of RNA degradation (11).

Some previous studies have indicated that RNAi components like Dicer play a role in tDR biogenesis (18-20), but our results suggest that *A. fumigatus* Dicer-like proteins play only a minor role. As compared to wild type, we did observe a decrease in the production of some tDRs in the conidia of Δ*ppdA*/B and an increase at later growth stages (i.e., 48-h-old hyphae). We also noticed alterations to tDR biogenesis in our inverted-repeat transgene overexpression strains, hinting at a more complex interplay between RNAi and tDR biogenesis in *A. fumigatus*. Argonaute proteins are known to stabilize tRFs in some systems (21), and it remains possible that the total pool of mature tRNAs susceptible to cleavage is lower in the Δ*ppdA*/B strain, based on the downregulation of genes associated with transcription observed in previous work (9). The mechanism of tDR biogenesis in *A. fumigatus* is unclear but we suspect that an Rny1 ortholog is likely involved (28, 45). Further study will be required to assign the enzymes responsible for fragment generation.

To better assess the tDR repertoire of *A. fumigatus*, we performed tDR-seq (46), which revealed an enriched pool of tRHs compared to sRNA-seq consistent with the literature (11). The tDR-seq dataset again revealed an abundance of rRNA reads, a smaller fraction of tDRs, and other RNAs consistent with previous work (47). In agreement with the sRNA-seq, a surprising absence of detection of tRF-3 CCA fragments derived from mitochondria-encoded tRNAs was noted. An improved annotation of the *A. fumigatus* mitochondrial genome and further molecular work will be required to better assess these missing CCA-containing fragments. The overall patterns of tDRs observed in our sRNA-seq was similar to that of the tDR-seq experiment, indicating some value remains in assessing tDRs in standard sRNA-seq for *A. fumigatus* as a first hint.

We took advantage of the limited endogenous sRNA landscape to assess the sRNAs produced by *A. fumigatus* upon induction of an inverted-repeat transgene. We were not surprised to observe an accumulation of 5’U-containing 20-nt sRNAs derived from the transgene, but it was notable that these sRNAs were primarily generated from two consecutive loci in the 3’ half of the 500-bp dsRNA. We also noted that sRNAs were produced in the wild-type and Δ*rrpA*/B strains, but not the Δ*dclA*/B and Δ*ppdA*/B strains, suggesting that the Argonaute proteins are required for either sRNA stability or biogenesis. We next assessed the ability of overexpression of the inverted-repeat transgene to induce growth arrest in liquid culture as previously observed in (9), but in the absence of RNAi components. We were again surprised to see that growth was limited in all cases without induction, even when the RNAi system was disrupted by deletion of RNAi components. The largest affect was observed for the Δ*rrpA*/B knockout strain after induction. It seems plausible that deletion of the RdRPs results in skewing of RNAi pathways towards more potent silencing via the canonical RNAi pathway, a phenotype consistent with the observation of an inhibitory role for the RdRP-dependent epimutation pathway on canonical RNAi in Mucor circinelloides (48). The growth disruption in Δ*dclA*/B and Δ*ppdA*/B strains lacking sRNAs (**Fig. 4A**) suggested that off-target effects of the sRNAs are not the only driver of growth arrest, but instead that excessive dsRNA itself may also serve as a signal for growth arrest. We hypothesize that in this case, high levels of dsRNA may compete with dsRBPs for endogenous dsRNA targets to impede normal biology, akin to a mycovirus infection (**Fig. 4E**). Such a mechanism would offer a natural brake on growth in response to abnormally high levels of dsRNA as during a mycovirus infection and likely give an advantage to uninfected cells.

The idea that dsRBP competition can influence function is not a new one (2, 49), and examples are found in the literature. For example, the human innate immune dsRBP sensor protein, protein kinase RNA-activated (PKR), was shown to form heterodimers that compete for binding to dsRNA to control regulation (50). The same receptor is also in competition with viral proteins like the vaccinia virus dsRBP E3L, which binds dsRNA to limit the activation of PKR (51). Similarly, an RNA-editing inactive version of the ADR1 enzyme of C. elegans, a human <u>a</u>denosine <u>d</u>eaminase <u>a</u>cting on <u>R</u>NA (ADAR) ortholog, was shown to control ADR-2 editing activity by outcompeting ADR-2 for binding sites (52). In plants, competition between dsRBPs was hypothesized to influence the selection of antiviral pathways upon infection with potato virus X (53), implicating cross talk between antiviral pathways during infection. In fungi, the details are less clear, but RNAi surely holds antiviral potential (10), and deletion of core machinery can lead to severe growth defects upon infection with mycoviruses (31). Future work will be required to determine the specific endogenous substrates that are outcompeted by exogenous dsRNA (e.g., during mycovirus infection or expression of an inverted-repeat transgene dsRNA).

In conclusion, our study provides 1) an improved view of tDR abundance and dynamics in *A. fumigatus* conidia and mycelium, which will likely be valuable in further defining the molecular biology of this important human pathogen; and 2) an interesting link between dsRNA metabolism and growth arrest in *A. fumigatus*, hinting at a potential growth checkpoint mediated by dsRNA in fungi that should be investigated further.

## MATERIALS AND METHODS

### Strains, culture conditions, and growth assays

Throughout this study, *A. fumigatus* strains (**Table S1**) were cultivated on Aspergillus minimal medium (AMM) agar plates and liquid culture as described previously (9). For measuring the dry weight, *A. fumigatus* strains were cultivated on AMM-agar plates for five days, then 1×10^8^ spores were added to 50-ml AMM liquid culture. Strains with inverted repeat elements were supplemented with 10 µg/ml of doxycycline. Flasks with AMM liquid culture were shaken at 200 rpm at 37°C in the dark for 24 h. Mycelium was collected through Miracloth (Millipore) and dried in the oven at 65°C for seven days prior to measurement of the dry weight. For the sRNA-seq of inverted-repeat transgene strains, 1×10^8^ freshly harvested spores were inoculated into 50 mL AMM, and incubated at 37°C and 200 rpm. After 16 h, 10 μg/mL of doxycycline was added to the inverted-repeat transgene strains (excluding wild type), followed by 8 h of additional growth (total of 24 h).

### RNA isolation for sRNA- and tDR-seq

RNA isolation was performed as described previously (9). Briefly, fungal mycelium was harvested from liquid culture using Miracloth (Millipore) and then disrupted in liquid nitrogen with a precooled mortar and pestle. Approximately 0.5 g of homogenized mycelium was transferred to a 2-ml Eppendorf tube. To this, 800 µl of TRIzol was added and vigorously vortexed. The tubes were briefly frozen in liquid nitrogen and then allowed to thaw on ice. Following thawing, 160 µl of chloroform was added to the mixture, vortexed, and centrifuged for 5 min at 4°C at maximum speed. The resulting aqueous upper phase was carefully transferred to a fresh 2-ml tube without disturbing the interphase. RNA extraction from the aqueous phase involved the addition of 1 volume of phenol/chloroform/isoamyl alcohol (25:24:1, v/v), followed by brief vortexing and centrifugation for 5 min at 4°C. This extraction process was repeated until no interphase was visible, and an additional extraction with 400 µl chloroform was performed. RNA was precipitated by adding 400 µl isopropanol for 20 min, followed by centrifugation for 20 min at 4°C. The resulting pellet was washed with 700 µl 70% ethanol and air-dried at 37°C for 5 min before being resuspended in RNase-free water. To remove any remaining DNA, a DNase treatment was carried out using 2 units of TURBO DNase (Thermo Fisher) per 10 µg RNA for 30 min at 37°C in a total volume of 100 µl. Finally, total RNA was collected using the RNA Clean and Concentrator-25 kit (Zymo Research) following the manufacturer’s instructions. DNase-free RNA concentrations were measured with QubitTM RNA BR Assay kit (Thermo Fisher Scientific, Darmstadt, Germany). Purified RNA was quantified using the QubitTM RNA BR Assay kit (Thermo Fisher Scientific, Darmstadt, Germany) on the Qubit Flex Fluorometer.

### RNA Isolation for IRT-sRNA-seq

For IRT-sRNA-seq, RNA was isolated generally as above with several improvements. Collected mycelium was first transferred to a 2-mL screw-cap tube filled 1/3 full with 0.5 mm soda lime glass beads (BoSpec). 1 mL of TRIzol (Thermo Fischer Scientific, Dreieich, Germany) was added to efficiently lyse the cells using the FastPrep-24™ Homogenizer by subjecting the cells to three rounds of 30 s, 4.0 m/s homogenization (1 min break between repetitions) and 5 min of incubation at room temperature. 200 μL of chloroform was added to each sample, vigorously mixed and incubated for 3 min at room temperature before centrifuging the samples at 12,000 x g for 15 min at 4°C. The upper aqueous phase containing the RNA was carefully transferred to a clean RNase-free tube and an equal volume of pure ice-cold ethanol was added and mixing thoroughly. Purification of RNA was carried out by using the RNA Clean and Concentrator™ kit (Zymo Research GmbH, Freiburg, Germany) following the manufacturer’s instructions. To eliminate DNase contamination, total RNA was treated with DNase I, RNase-free (Thermo Fisher Scientific, Darmstadt, Germany) at 1 U per μg of RNA. Following a 1-h incubation at 37°C, the reaction was terminated by adding 2 μL of 50 mM EDTA at 65°C for 10 min and processed further as described above.

### Library preparation and sequencing

sRNA-seq libraries were prepared by Novogene using the TruSeq Small RNA Library Preparation Kit according to the manufacturer’s protocol, then sequenced using NovaSeq 6000 instrument in a single-read, 50-bp based mode at 10 million reads of sequencing depth. For tDR-seq performed by CD Genomics (tRF&tiRNA-seq), the following treatments for modifications were performed: 1) 3’-aminoacyl (charged) deacylation to 3’-OH for 3’ adaptor ligation, 2) 3’-cP removal to 3’-OH for 3’ adaptor ligation, 3) 5’-OH (hydroxyl group) phosphorylation to 5’-P for 5’-adaptor ligation, and 4) m^1^A and m^3^C demethylation for efficient reverse transcription. Finally, RNA libraries were sequenced using an Illumina HiSeq 2500 instrument in a 50-bp single-read mode at 10 million reads of sequencing depth.

### Data Analysis

TrimGalore version 0.6.8 (https://github.com/FelixKrueger/TrimGalore) was used for quality checks and removing adaptors of sequencing reads. SortMeRNA version 4.3.7 (54) was used to remove potential rRNA reads, with the rRNA database generated from Rfam (https://rfam.org/) and silva (https://www.arb-silva.de/). In silico removal of rRNA reads did not alter analysis results and was subsequently omitted for downstream analyses. Sequencing data were analyzed with unitas v. 1.7.0 (55) using default parameters and *Aspergillus fumigatus* A1163 ASM15014v1 as reference genome. *A. fumigatus* nuclear tRNAs were referenced from GtRNAdb (56), mitochondrial tRNAs were scanned from the mitochondrial genome (57) using the 27 tRNAs predicted by tRNAscan-SE (58). All analyses were performed using “R” version 4.3.1 (2023-06-16) and Rstudio (2022-02-02), an integrated development environment (IDE) for R or Prism 10.4.0 software (GraphPad Software). Figures were drawn with the ggplot2 package (version 3.4.2) in R. Graphs for read coverage were created using the SparK package (59). P-values were determined by ANOVA or two-tailed Student’s t-tests where appropriate and differences between the groups were considered significant at a P-value of <0.05.

## Data availability

The sRNA-seq data are available under NCBI Gene Expression Omnibus (GEO) identifier GSE267454. These data were collected simultaneously with previously published mRNA-seq data (9) available using identifier GSE223618. The tDR-seq data is available under identifier GSE267453. The sRNA-seq for the inverted-repeat transgene overexpression is available using GSE286229.

## ACKNOWLEDGEMENTS

We would like to thank Pamela Lehenberger and Bhawana Israni for excellent technical assistance and Axel Brakhage, Alexander Bruch, and Xiuqiang Chen for helpful discussions. The work presented here was generously supported by the Federal Ministry for Education and Research (BMBF: https://www.bmbf.de/), Germany, Project FKZ 01K12012 “RFIN – RNA-Biologie von Pilzinfektionen”. Additional funding support came from the Deutsche Forschungsgemeinschaft (DFG; German Research Foundation) under Germany’s Excellence Strategy – EXC 2051 – Project-ID 390713860. The funders had no role in the design of the study. The authors declare no conflicts of interest.

## Notes

### Competing Interest Statement

The authors have declared no competing interest.

### Summary of Updates

The revised version contains improved data analysis for most of the prior analyses and several additional experiments assessing the production of small RNAs in response to overexpression of an inverted-repeat transgene. The manuscript is written with a new framework to hihglight additional relevance to the community. Two additional authors are added for their support on new experiments.

