## Supplemental Information for "Investigation of *Aspergillus fumigatus* small RNA biogenesis uncovers evidence of double-stranded RNA-dependent growth arrest"

**Running Title:** Excess dsRNA limits *A. fumigatus* growth.

**Key Words:** filamentous fungi, fungal pathogen, tRNA-derived RNAs, small RNA, double-stranded RNA

**SUPPLEMENTARY FIGURES AND LEGENDS**

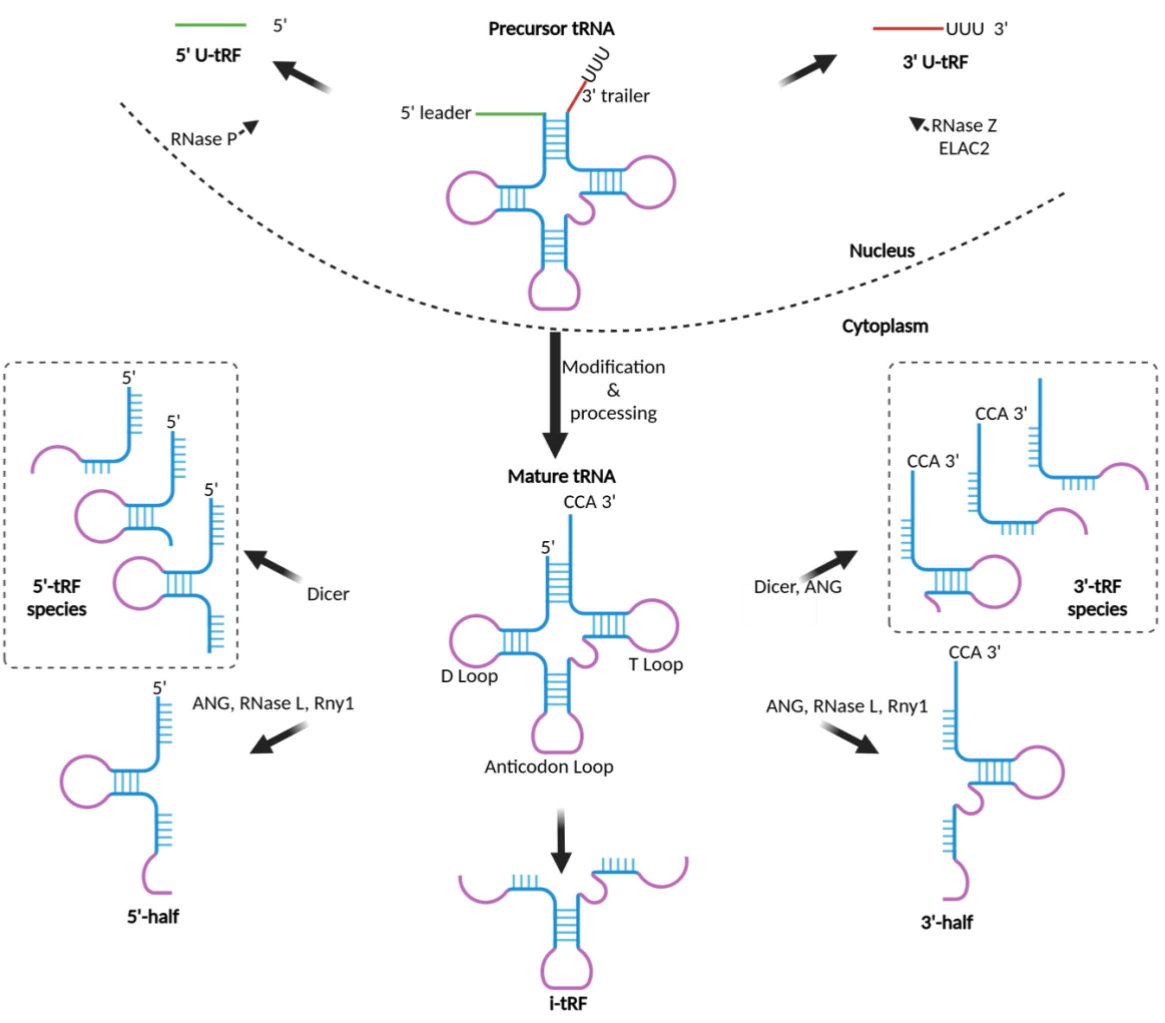


**FIG S1. Model of putative tDR biogenesis.**

A schematic showing the relevant tRNA-derived RNAs (tDRs) discussed in this manuscript based on the literature [1, 2]. tDR = tRNA-derived RNA; tRF = tRNA fragments; tRF-3 = 3' fragments; tRF-3 CCA = tRF-3 fragments including CCA tail; tRF-5 = the 5' fragments; tRH = tRNA halves; 5'tRH = 5' tRNA half; 3'tRH = 3' tRNA half. Created in BioRender. Blango, M. (2025) https://BioRender.com/i00l091.

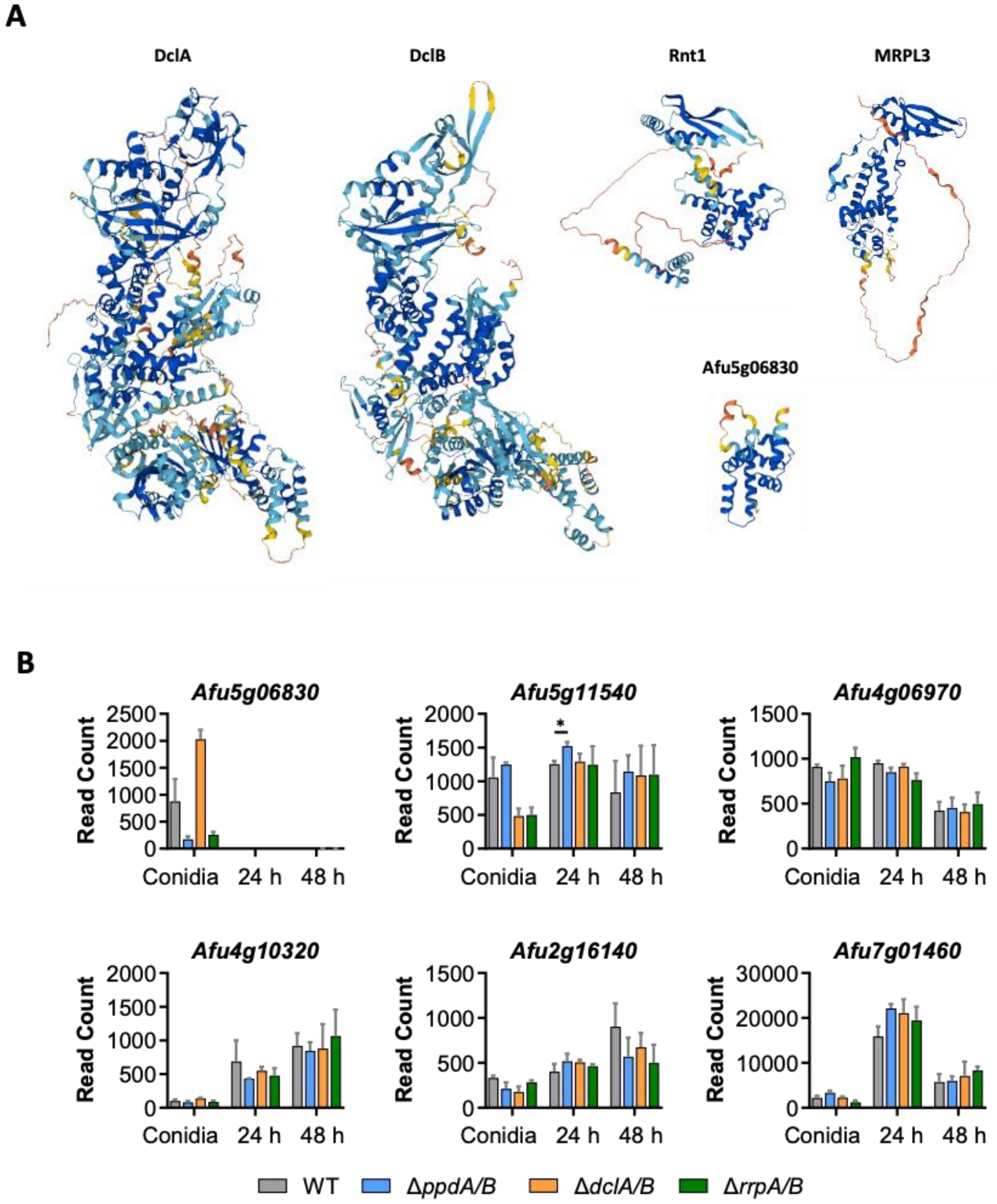


**FIG S2.** ***A. fumigatus*** **harbors ten annotated dsRBPs with five containing putative RNase III domains.**

**(A)** Alphafold structure predictions for the five annotated RNase III-domain containing proteins of *A. fumigatus* (Uniprot). **(B)** Median ratio normalization (MRN) expression of the remaining dsRBP genes not previously shown in (Figure 5 of [3]) in conidia and mycelium. Values are indicated as read count of individual genes in conidia, 24-h and 48-h mycelium using data from mRNA-seq experiment with NCBI Gene Expression Omnibus (GEO) identifier GSE223618.

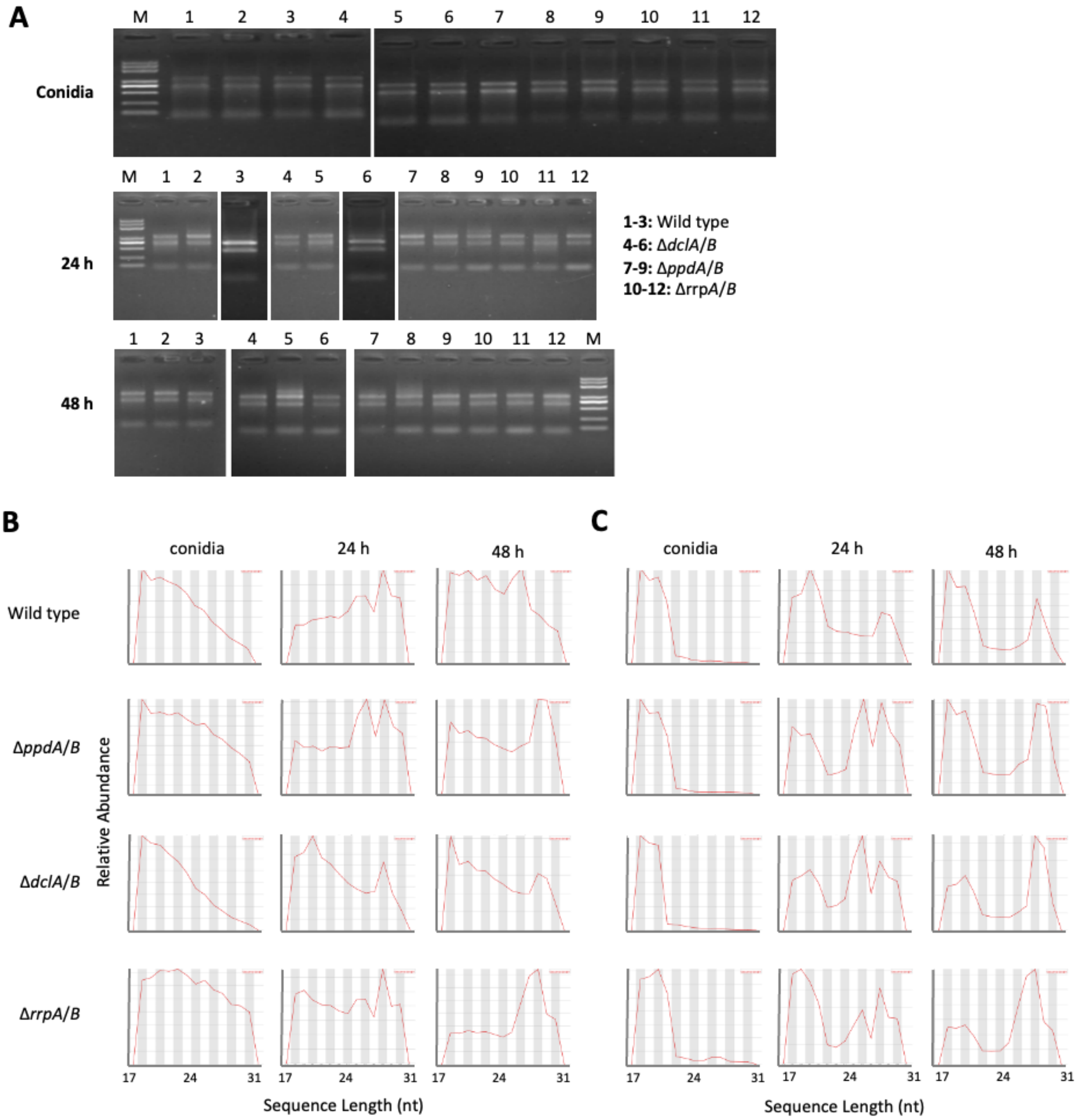


**FIG S3. RNA quality and read processing for sRNA-seq.**

**(A)** Gel image from denaturing agarose gel electrophoresis of isolated total RNA from *A. fumigatus* conidia and mycelium for sRNA-seq (Novogene). Each condition consists of three separate biological replicates. Read distribution plots for cleaned up reads **(B)** and reads after additional rRNA removal **(C)** created using FastQC. SortMeRNA version 4.3.7 [4] was used to remove potential rRNA reads, using an rRNA database generated from Rfam (https://rfam.org/) and silva (https://www.arb-silva.de/).

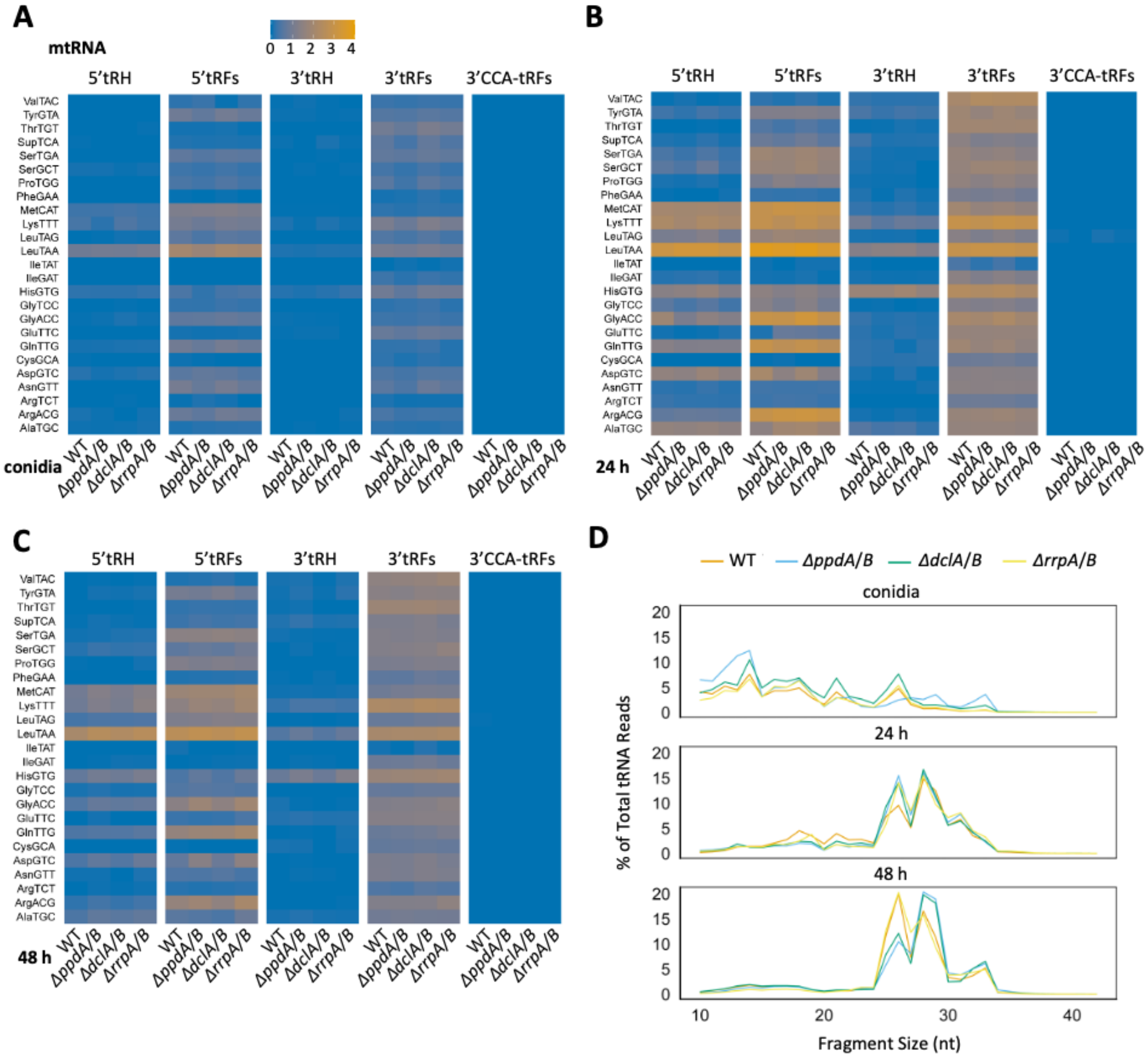


**FIG S4. Heatmap showing expression of mitochondria-encoded tDRs.**

Heatmap showing abundance of mitochondria-encoded (mt) tDR species and their tRNA origin across sampled morphotypes (Conidia **(A)**, 24 hours- **(B)** and 48 hours- **(C)** old hyphae) of wild-type and RNAi double knockout strains. Plots show counts per million mapped reads based on fractionated counts produced by unitas v. 1.7.0 [5]. n = 3 biological replicates per condition. **(D)** Read length distribution for tDRs of mitochondria-encoded tRNAs from conidia and mycelium (24-h and 48-h liquid cultures).

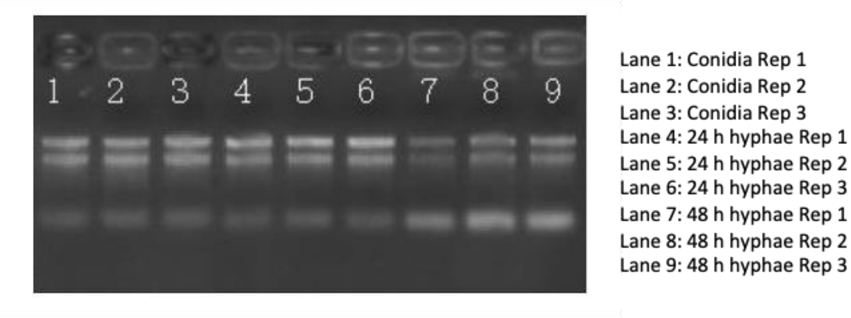


**FIG S5. Total RNA input quality for tDR-seq.**

Gel image from denaturing agarose gel electrophoresis of isolated total RNA from *A. fumigatus* conidia and mycelium for tDR-seq (CD Genomics). Each condition consists of three separate biological replicates.

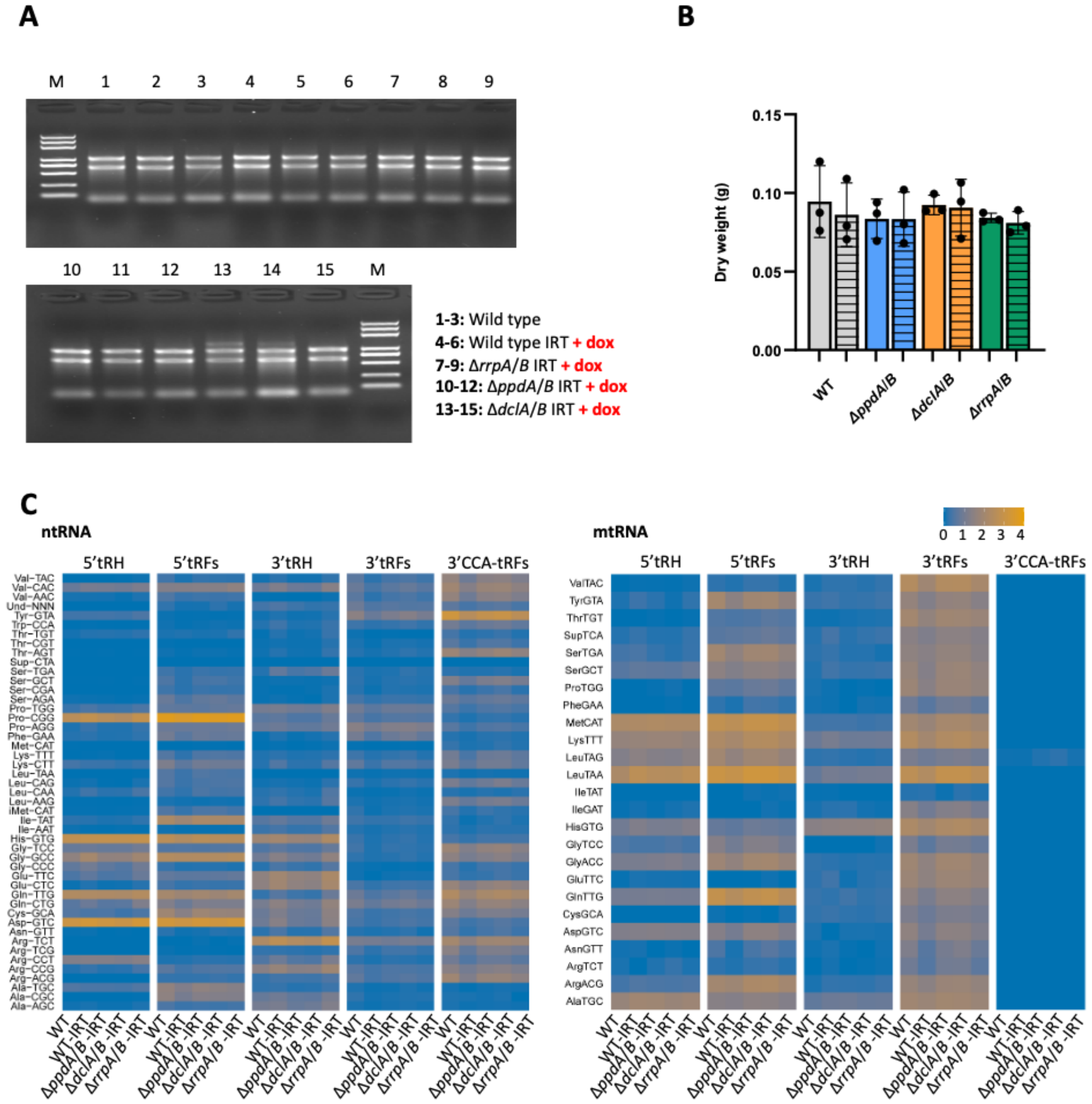


**FIG S6. Expression of inverted-repeat transgene induces minor alterations to tDR profile.**

**(A)** Gel image from denaturing agarose gel electrophoresis of isolated total RNA from *A. fumigatus* mycelium for sRNA-seq (Novogene). Each condition consists of three separate biological replicates. **(B)** Measurement of fungal dry weight (biomass) after 24-h growth in liquid culture in the presence or absence of doxycycline for the noted RNAi knockout strains or compared to a wild-type strain. Relevant comparisons were performed with two-way ANOVA and Šídák's multiple comparisons posttest from three biological replicates, with no $P$-values < 0.05 identified. **(C)** Heatmap showing the abundance as determined by sRNA-seq of tDR species from nuclear- (left) and mitochondria-encoded tRNA (right) (tRNA Transcript ID =1, Gene locus ID =1) of 24 h-old-mycelium of wild-type or RNAi double knockouts expressing a *pksP* inverted-repeat transgene as noted. Plots show counts per million mapped reads based on fractionated counts produced with unitas v. 1.7.0 [5]. n = 3 biological replicates per condition.

### SUPPLEMENTARY TABLES

**Table S1.** List of strains used in this study.

| **Strain** | **Genotype and/or phenotype** | **Reference** |
|---|---|---|
| ***A. fumigatus*** | | |
| CEA17Δ*akuB* | *akuB*$^{KU80}$::*pyrG*; PyrG$^{+}$, Δ*akuB* | [6] |
| Δ*dclA* Δ*dclB* | CEA17Δ*akuB*, PtrA$^{R}$, Hyg$^{R}$ | [3] |
| Δ*ppdA* Δ*ppdB* | CEA17Δ*akuB*, Phleo$^{R}$, PtrA$^{R}$ | [3] |
| Δ*rrpA* Δ*rrpB* | CEA17Δ*akuB*, Hyg$^{R}$, PtrA$^{R}$ | [3] |
| CEA17Δ*akuB_pksP*-IRT | CEA17Δ*akuB*, Hyg$^{R}$ | [3] |
| Δ*dclA* Δ*dclB _pksP*-IRT | CEA17Δ*akuB*, PtrA$^{R}$, Hyg$^{R}$, Phleo$^{R}$ | [3] |
| Δ*ppdA* Δ*ppdB B_pksP*-IRT | CEA17Δ*akuB*, Phleo$^{R}$, PtrA$^{R}$, Hyg$^{R}$ | [3] |
| Δ*rrpA* Δ*rrpB _pksP*-IRT | CEA17Δ*akuB*, Hyg$^{R}$, PtrA$^{R}$, Phleo$^{R}$ | [3] |

### SUPPLEMENTARY DATASETS

**Dataset S1.** Normalized abundance from unitas analysis of sRNA-seq on wild-type and RNAi double knockout strains.

**Dataset S2.** Output of Manatee software using sRNA-seq on wild-type and RNAi double knockout strains as input.

**Dataset S3.** Normalized abundance from unitas analysis of tDR-seq on wild-type *A. fumigatus*.

**Dataset S4.** Normalized abundance from unitas analysis of inverted-repeat transgene sRNA-seq.
